# Evolution of Animal Neural Systems

**DOI:** 10.1101/116715

**Authors:** Benjamin J. Liebeskind, Hans A. Hofmann, David M. Hillis, Harold H. Zakon

## Abstract

Nervous systems are one of the most spectacular products of evolution. Their provenance and evolution have been an area of interest and often intense debate since the late 19th century. The genomics era has provided researchers with a new set of tools with which to study the early evolution of neurons, and recent progress on the molecular evolution of the first neurons has been both exciting and frustrating. It has become increasingly obvious that genomic data is often insufficient to reconstruct complex phenotypes in deep evolutionary time. We review this recent progress and its attendant challenges, and suggest ways forward.

## 1. INTRODUCTION

Behavior is widespread on the *unicellular* tree of life. When Anton van Leeuwenhoek looked through his microscope, saw swimming microorganisms, and called them animalcules due to their rapid motions, he was the first witness to the remarkable similarities between the behavior of microbes and animals [1]. But among multicellular organisms, animals are preeminent in the rapidity and diversity of their behavioral repertoire. This is due to key differences in the ways that animals achieved a multicellular lifestyle as compared to plants, fungi, and red and brown algae, the other major multicellular eukaryotic lineages. Development of novel pathways for cell adhesion [2] and intercellular signaling [3] laid the groundwork, but the advent of neurons and muscles were the key adaptations that allowed animals to overcome the behavioral bottleneck that attended multicellularity in every other eukaryotic lineage.

With the sequencing of animal genomes from diverse phyla [4-7] and an emerging consensus over the broad animal phylogeny [8], a lively debate has sprung up over the nature of the early evolution of neurons, and, more broadly, how to interpret genomic data in a way that best enlightens the deep origins of complex tissues types [4,9-12]. In the last two years alone, there have been five journal issues dedicated completely or in part to the early evolution of nervous systems. This debate is a continuation of a perennial debate over the nature of the first neurons [13,14], but for the first time, the debate is centered on the interpretation of molecular and genomic, rather than phenotypic or physiological, data. At the end of the last era of comparative nervous system research, George Mackie remarked that the distinction between neurones [*sic*] and non-nervous cells has become somewhat blurred and it now seems most appropriate to ask not which cell lineages originally gave rise to nerves but where the genes expressed in neurogenesis originally came from [14]. We have made significant progress towards this goal, but what have we learned about cell-level phenotypes? As might be expected, molecular data cannot tell us everything we want to know. Below, we give a broad overview of recent work on the molecular evolution of the first neurons, highlight its limitations, and discuss areas for future research.

## 2. BACKGROUND

### 2.1. What is a nervous system?

By nervous system we typically mean the network of neurons that underlie animal behavior. It has long been appreciated that nervous system is an imprecise term [13]. Many other cell types beside neurons are nervous, i.e. electrically excitable, and exist in systems, such as pancreatic or muscle cells. Plants and unicellular organisms also make use of electrical excitability to mediate behavior (see below), but are not said to have nervous systems. It is the presence of neurons that really distinguishes animal nervous systems, so we prefer the term neural systems. Neurons are typically defined morphologically and physiologically as an elongated and excitable cell that makes a synapse onto another cell. To this common and admittedly vague definition (glia cells can also be excitable and make synapses), we add the functional qualification that neurons are usually said to encode information. That is, their particular activity is an arbitrary symbol or code of downstream activity, ultimately, of animal behavior. A neural system encodes information in two ways: First, in a dynamic electrical code within neurons, and second, in the wiring code or connectome between neurons. Both within- and between-cell signaling can be dynamically altered, and this plasticity is responsible for learning, memory, development, and behavioral complexity.

### 2.2. Phylogenetic context of early neural evolution

When did the first neurons arise, and how? Although this question is still hotly debated, the phylogenetic context for the early neural evolution is starting to come into light as a revised animal tree of life has emerged. One striking development has been the realization that many traits considered to be synapomorphies of major animal clades, such as a through-gut, epithelia, and neurons, are not in fact synapomorphic [8]. Perhaps the most radical change to the animal tree of life has been the growing consensus that ctenophores, which have a complex neural system and behavior, branched off first from the remaining animal lineages (Figure 1). Sponges had previously been considered to be the earliest branching lineage and their lack of neurons and sedentary lifestyle (as adults) was thought to be the ancestral condition of animals. Though it is still possible that the placement of ctenophores results from problems in phylogenetic inference [15], the early splitting of ctenophores is supported by a variety of careful studies [4,5,16-18]. This placement of ctenophores raised the question of whether neurons arose once in animal evolution and were lost in sponges and placozoans (Figure 1), or arose independently in ctenophores and Planulozoa (cnidarians+bilaterians) [4,17].

**Figure 1.**
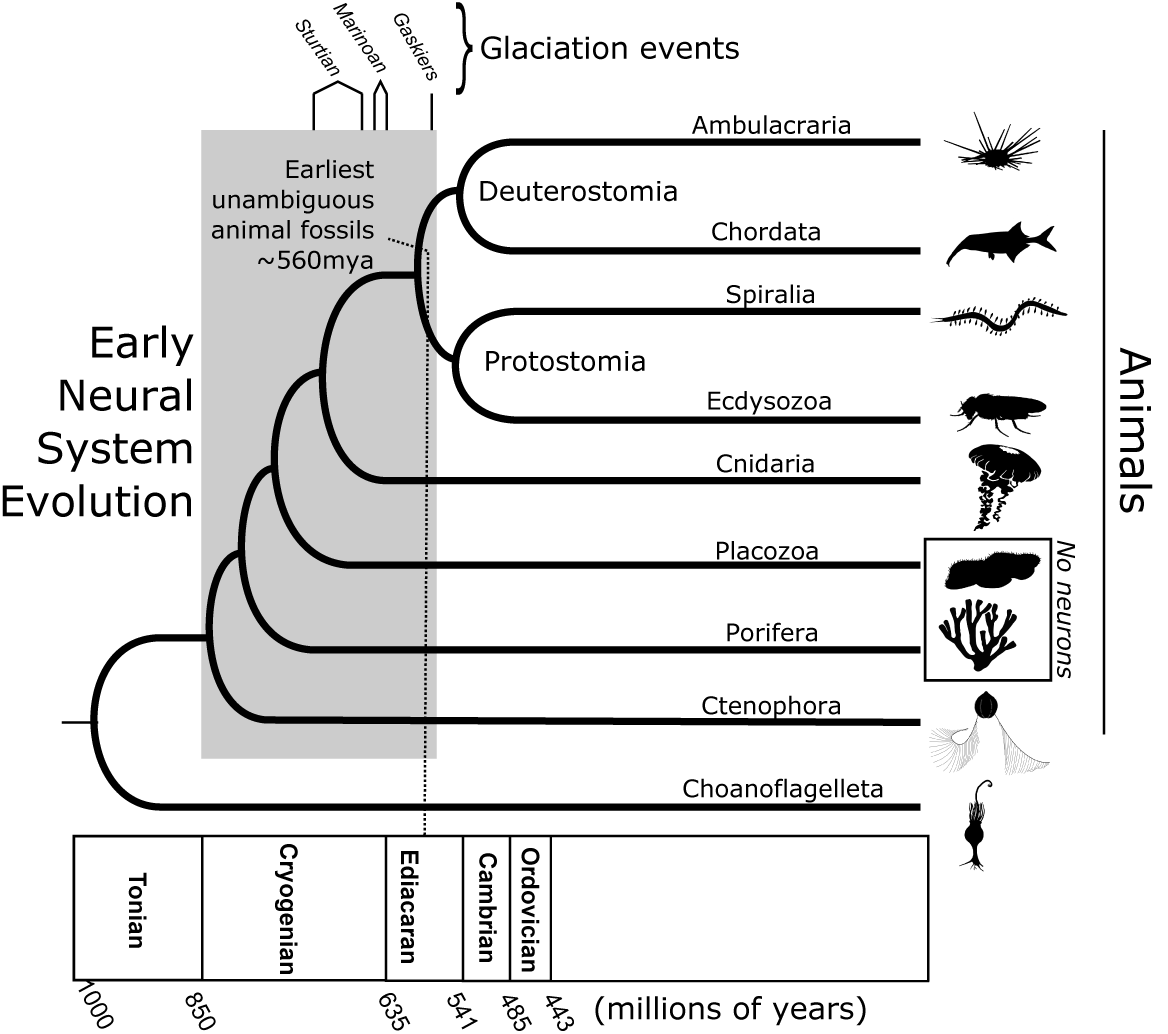
The deepest animal divergences pre-date unambiguous animal fossils by hundreds of millions of year, even taking plausible confidence intervals into account [21]. The most likely explanation for this discrepancy is that the first animals were too small and simple to fossilize as recognizable metazoans. Early neurons arose during this period, so their origins must be reconstructed primarily from extant animal lineages. Silhouettes here and throughout are from Phylopic (http://phylopic.org/).

The placement of sponges and ctenophores is important, but not decisive, for reconstructing the origin(s) of neurons, however [19]. The debate over the tree topology has often assumed that the two scenarios mean that the common animal ancestor was either ctenophore-like or sponge-like, but neither is likely to be the case. The deepest divergences of extant animal lineages stretch back to the Cryogenian, long before the appearance of large animal fossils (Figure 1) [20,21]. It is thus entirely plausible that animals began diversifying before becoming more complex and that these early animals were too small and delicate to fossilize or to be recognizable as animals. Importantly, no extant lineage is frozen in time, so resolution of the tree is necessary, but not sufficient for ancestral character recon-struction. The early evolution of animal neural systems occurred during this dark period in the Cryogenian (Figure 1), so we must use extant molecular and physiological data to ask whether neurons evolved once, or repeatedly.

### 2.3. The promise and perils of genomic data

Despite all the new genomic data, the field has failed to reach an agreement on the nature of the early evolution of neurons. Why have we failed to do so? It is simply because we cannot perfectly predict phenotypes from genomic data alone. Why can’t we? Because the structure of proteins and molecular systems is not uniquely determined by the phenotypes they encode, and vice versa. There is a many-to-many mapping between mechanisms and phenotypes. For instance, homologous phenotypes can diverge in molecular composition (systems drift; [22]), and homoplastic phenotypes can independently recruit homologous molecular modules (deep homology; [23]), so molecular similarities and differences are not necessarily reflective of homology or homoplasy at the phenotypic level. Because complex cell types will contain proteins with a wide diversity of evolutionary histories, one can always find molecular evidence to support ones hypothesis of homology or of homoplasy. Correspondingly, there are no universal molecular markers of neurons [24], and even if there were it would not rule out the possibility of homoplasy. A robust determination of homology must rely on a better understanding of how cell level phenotypes co-evolve with molecular modules than is currently available. Thus, the next challenge to the field is to start putting the genomic data back into its physiological context, and this will involve developing comparative methods that explicitly take hierarchical systems into account.

Because neurons are among the most complex and best studied cell types, comparative neuroscience offers an ideal system for hierarchical comparative methods. Neurons, and still more, neural systems, are not monolithic entities. They consist of numerous molecular machines and their constituent proteins, each carrying out the processes we identify with neural function (Figure 2). Below, we review these molecular systems and recent advances in our understanding of how they evolved in the first animals, highlighting areas where little is known. Finally, we end with an example of how genomic data must be contextualized to be interpreted, by exploring the surprisingly plastic evolution of the neuromuscular junction.

**Figure 2.**
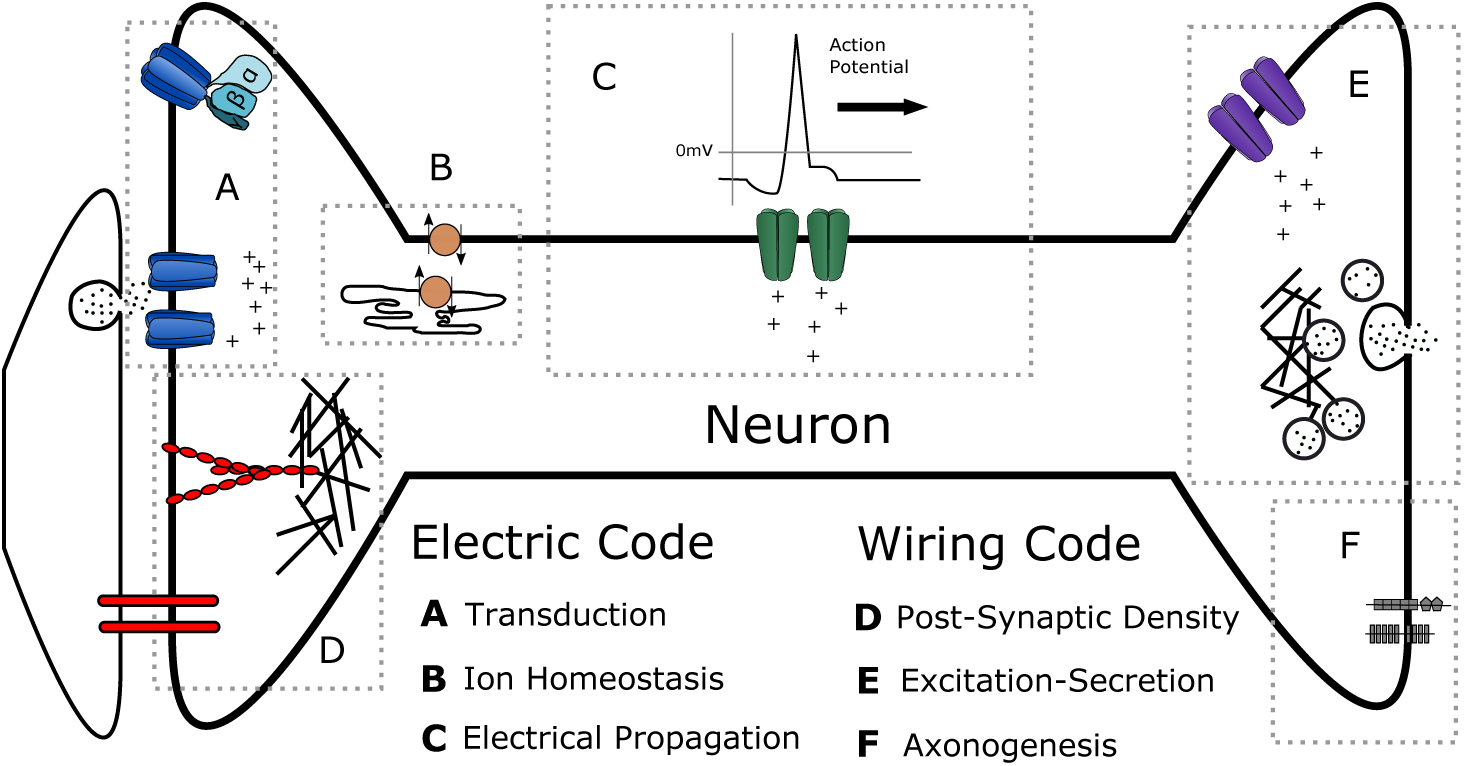
The molecular modules of a mature neuron can be divided into modules underlying the electrical code within neurons, and those underlying the synaptic code between neurons. Each of these two sections can be further divided into three sub-modules. Proteins from all modules were present in the unicellular ancestor of animals.

## 3. ELECTRICAL CODE

Electrical excitability is a defining feature of neurons. Most neurons propagate all-or-none electrical disturbances called action potentials. These spikes can be sent over meters-long axons and arrive at the nerve terminal with near-perfect fidelity, and thus constitute a digital signal. Besides fidelity, conduction speed is a major adaptive feature of neurons, allowing organisms to move rapidly. Thus, giant axons, which reduce internal resistance, have evolved multiple times as part of startle-escape behavior circuits, most famously in crayfish, squid, and teleost fish, but also in cnidarians and ctenophores [25-27]. Myelination, which insulates axons, was previously thought to be a vertebrate-specific innovation, but in fact has evolved several times in protostomes as well [28]. The creation of the neural electrical code relies on three modules of proteins: one that creates the potential energy for the action potential (ion homeostasis), one that transduces sensory and intercellular signals into the electrical code (transduction), and one that propagates the electrical signals along neurons (propagation) (Figure 2).

### 3.1. Ion homeostasis

The flow of ions that creates an action potential is powered by actively maintained electrochemical gradients. In this, neurons resemble all other cell types, which must also regulate the distribution of different ions across their membrane. In neurons, the key tasks of ion homeostasis are maintaining a low cytoplasmic calcium level to protect the fidelity of intracellular calcium signaling and maintaining a negative voltage across the membrane to power action potentials.

The primary means of maintaining membrane potential is active pumping via ATPase proteins, which in humans use up to 10% of the bodies at rest in the brain alone [29]. P-type (for phosphorylated) ATPases are the most important for neuronal function, and can be found in all cellular life forms [30]. Na^+^/K^+^ ATPases can be found in the genomes of many non-animals, where they are presumably used for maintaining ion homeostasis in high-salt environments [31]. Proteins that maintain low cytoplasmic calcium levels, such as the sarco-/endoplasmic reticulum calcium pump (SERCA) and PMCA pumps, Na^+^ /Ca^2+^ exchangers (NCX), and mitochondrial calcium uniporter (MCU), are found across eukaryotes as part of a pan-eukaryotic calcium signaling toolkit [32,33].

The origin of neurons does not seem to have relied on an increased number of genes for these pumps and exchangers. For instance, humans have 24 P-type ATPases, whereas yeast have 14 and flies 12. The plant *Arabidopsis thaliana* has 46 [30]. Thus, the ion homeostasis module is both evolutionarily ancient, and pleiotropically expressed in multiple cell types. Though there may have been protein-level specializations associated with the origin of neurons in this module, they are not currently well-understood.

### 3.2. Transduction

The first step of neural signaling is often the transduction of an input signal, either from the environment or another cell, into an electrical signal. This occurs both at synapses between neurons and at sensory neurons, such as photoreceptors or olfactory neurons. These two distinct tasks, sensory transduction and synaptic transduction, are often treated separately, but many of the same proteins are expressed in both sub-modules across animals. Of the many protein families involved in transduction, only the amiloride-sensitive channels (ASC, also called EnaC, ASIC, or DEG), and the Pannexin families are metazoan novelties all others were present in the unicellular ancestors of animals [5,34,35], where they were probably used for environmental sensing.

Two major families of proteins transduce environmental stimuli into electrical signals in vertebrate neurons: transient receptor potential (TRP) channels and G-protein coupled receptors (GPCRs) [36,37]. Whereas TRP channels transduce environmental signals directly to electrical signals, GPCRs operate indirectly on downstream ion channels via the G-protein signaling pathway. GPCRs form the largest gene super-family in tetrapods, with the majority being olfactory receptors [38]. Individual GPCR genes are rapidly duplicated, lost, or inactivated [39]. GPCRs also mediate taste and photoreception. This latter sensory modality is mediated by opsins, which bind a chromophore whose response to light sets off the G-protein signaling cascade. There have been independent expansions of opsins in cnidarians [40], notably in cubozoan jellyfish, which represent one of three lineages of animals to have evolved lens-bearing eyes [41]. Curiously, opsins were lost in the sponge lineage and yet ciliated sponge larvae are phototactic. This behavior is mediated by cryptochromes instead of opsins [42], probably a case of systems drift.

Whereas mammalian odorant receptors are all GPCRs, hexapod odorant receptors are all ion channels. Some belong to the same class of 7 transmembrane segment proteins as do GPCRs (odorant receptors [ORs] and gustatory receptors [GRs]; [37,43]), and others are part of the ionotropic glutamate receptor (iGluR) family (ionotropic receptors [IRs]; [44]), which in vertebrates are solely expressed in synapses. Nematodes have a large number of GPCRs, some of which play a role in chemoreception, as well as orthologs of IRs whose function is unclear [43]. Cnidarians have homologs of hexapod GRs that curiously play a role in development, with no indication of a role in chemosensation [45]. Thus it is not clear whether the bilaterian ancestor primarily used GPCRs for chemoreception, ion channels, or both. TRP channel families are not nearly as large as GPCRs, but are molecularly diverse. They underlie mechanosensation, thermosensation, and certain types of tastes, such as capsaicin and menthol [46]. The ASC family plays a role in both sensation (salt taste) synaptic transmission [47], with a larger role in synaptic transmission in cnidarians [48].

Like sensory receptors, synaptic channels and receptors exist in large gene families that have undergone substantial independent expansions in a number of animal lineages [35]. In vertebrates, the most important synaptic receptors are either GPCRs, or ion channels of the iGluRs or pentameric ligand gated ion channel families (pLGICs, also called Cys-loop receptors). Nearly all the major neurotransmitters target both ionotropic and metabotropic receptors, and there are key differences between non-vertebrate and vertebrate receptors that suggest widespread convergent evolution for ligand specificity, just as in olfactory sensation. For instance, ionotropic glutamate receptors in vertebrates are all part of the ion channel super-family iGluR, which are an ancient family of tetrameric channels distantly related to voltage-gated potassium channels [49]. These channels create an excitatory postsynaptic potential (EPSP) by letting in Na^+^ and Ca^2+^. In protostomes, such as *Drosophila* and *C. elegans*, however, some ionotropic glutamate receptors belong to the pentameric pLGIC channel family and create an inhibitory response by passing Cl^-^ anions. Moreover, these protostome glutamate receptors have evolved several times independently [50,51]. Ctenophores have lost pLGICs all together, the only lineage with a neural system known to have done so, and may rely on other channel families for the same function, such as iGluR and ASCs [4,5,35]. Supporting this hypothesis, it has recently been shown that the radiation of the iGluR family in ctenophores has evolved both glutamate and glycine-sensitive isoforms convergently with bilaterians [52].

Thus the proteins involved in the transduction of extracellular signals into electrical signals in neurons are characterized by large gene families with rapid turnover and many instances of convergent evolution of ligand specificity. At synapses, receptors may originate by first binding a molecule that is present as bi-product of biosynthesis, a process that has been called ligand exploitation [53]. So from a molecular evolution standpoint, transduction proteins are the opposite of the slowly evolving, highly pleiotropic ion homeostasis proteins.

### 3.3. Electrical propagation

Local electrical signals are transient and will attenuate over space. To propagate signals over large distance, neurons use ion channels that respond to voltage itself, called voltagegated ion channels. Voltage-gated ion channels and the regenerative potentials they create are found across the tree of life. For instance, plants with rapid behaviors, such as the Venus flytrap (*Dionaea*) or the sensitive plant (*Mimosa*), trigger these behaviors with ionic action potentials conducted through the phloem [54]. Protists also use action potentials to trigger bioluminescence [55], to coordinate cell deformations [56], and for control of ciliary beating [57]. Bacterial biofilms also use regenerative potential changes to coordinate colony growth [58]. In animals, neurons are not the only cell types to make use of electrical excitability: insulin release from the beta cells of the pancreas, muscle contraction, and chemotaxis in sperm are all mediated by electrical signaling [59,60].

In non-animals, the change in electric potential is often a byproduct of the main task of the impulse: the delivery of an ion species that effects a change in cell biology. Typically, this ion is calcium, which triggers a variety of cellular processes. In plants, the action potential creates an osmotic potential, and the resultant turgor pressure is the plant equivalent of muscle contraction. However, in animal axons, the action potential is a signal, and does not substantially alter the cell biology of an axon until it arrives at the terminus. The key change that allowed this development was the evolution of voltage-gated sodium (Na_*v*_) channels from pre-existing voltage-gated calcium channels (Ca_*v*_), as Na^+^ does not broadly trigger cellular signaling pathways as Ca^2+^ does. This allowed animal axons to generate action potentials at a high rate without poisoning the cell with Ca^2+^ [59].

After a positive displacement by Na_*v*_ channels, action potentials are then repolarized by K^+^ efflux through voltage-gated potassium channels (K_*v*_). K_*v*_ channels therefore play the central role in shaping the neural electric code and are a large and diverse family. While there are typically fewer than 5 Na_*v*_ genes in animal genomes (though they have radiated to 8 in fish and 10 in tetrapods), there are typically 10 times as many K_*v*_ genes [59]. Another member of the voltage-gated family has lost its sensitivity to voltage, and has been called sodium leak channel non-selective, or NALCN. This channel family keeps neurons near the threshold for Na_*v*_ activation and is crucial for rhythmically firing neurons that mediate, e.g., breathing in mice and snails [61].

Na_*v*_s evolved from Ca_*v*_s in a stepwise fashion, with bona fide Na^+^-selective channels arising twice, once in cnidarians, and once in the bilaterian ancestor [62,63], suggesting convergence toward a complex neural code in these lineages. NALCN channels have also likely converged towards Na^+^ permeability via similar mutations in their ion selectivity filter [64,65]. Thus sodium channels have arisen numerous times by convergent means from calcium channels.

In animals, the K_*v*_ gene family has radiated independently in a number of lineages [35,66], presumably for the regulation of an independently expanding electric code. In a remarkable case of convergence, Erg-type K_*v*_ channels have radiated independently in cnidarians and bilaterians, and in both radiations converged towards two distinct types of pore closure [67].

Thus the voltage-gated channels that mediate electrical signaling in neurons are characterized by convergent biophysical phenotypes, and the broad similarities between electrical signaling in cnidarians and bilaterians are largely a result of convergent evolution. Almost nothing is known about the physiology of ctenophore neurons, though their muscles make mostly calcium-based action potentials [68]. Glass sponges can propagate electrical impulses through their syncytial tissue [69], but demosponges (and perhaps other sponge lineages) have lost all voltage-gated channels genes [6,35]. Thus much remains to be discovered about the early evolution of electrical signaling.

## 4. WIRING CODE

Besides the electrical code within neurons, animal behavior is also encoded in specific wiring diagrams or connectomes between neurons. The connections between neurons, and between neurons and downstream effector cells, occur at specialized cell junctions called synapses. Synapses between neurons can either be (1) electrical, where the communication is direct ion flow through protein channels that connect the cytoplasm of the two cells; (2) chemical, where communication is via the release of a neurotransmitter and the cells are not directly coupled; or (3) mixed [70]. Though it has recently been shown that electrical synapses can undergo use-dependent plasticity [71], the mechanisms of long-term plasticity, which underlie learning and memory, are much better understood (and probably more important) at chemical synapses, which we focus on here. The proteins involved in chemical synaptic transmission are much more numerous and diverse than those involved in electrical conduction and function in medium-to-large protein complexes whose structure and function are not yet fully understood (Figure 3). In vertebrates, synaptic transmission is usually in one direction (as shown in Figure 3), but ctenophore and cnidarian synapses are often bidirectional [72,73]. Little is known about the proteins involved in cnidarian and ctenophore synapses, and there is probably much undiscovered complexity there. The wiring code can be divided into pre- and post-synaptic modules, and a module that specifies the wiring diagram during development (Figure 3).

**Figure 3.**
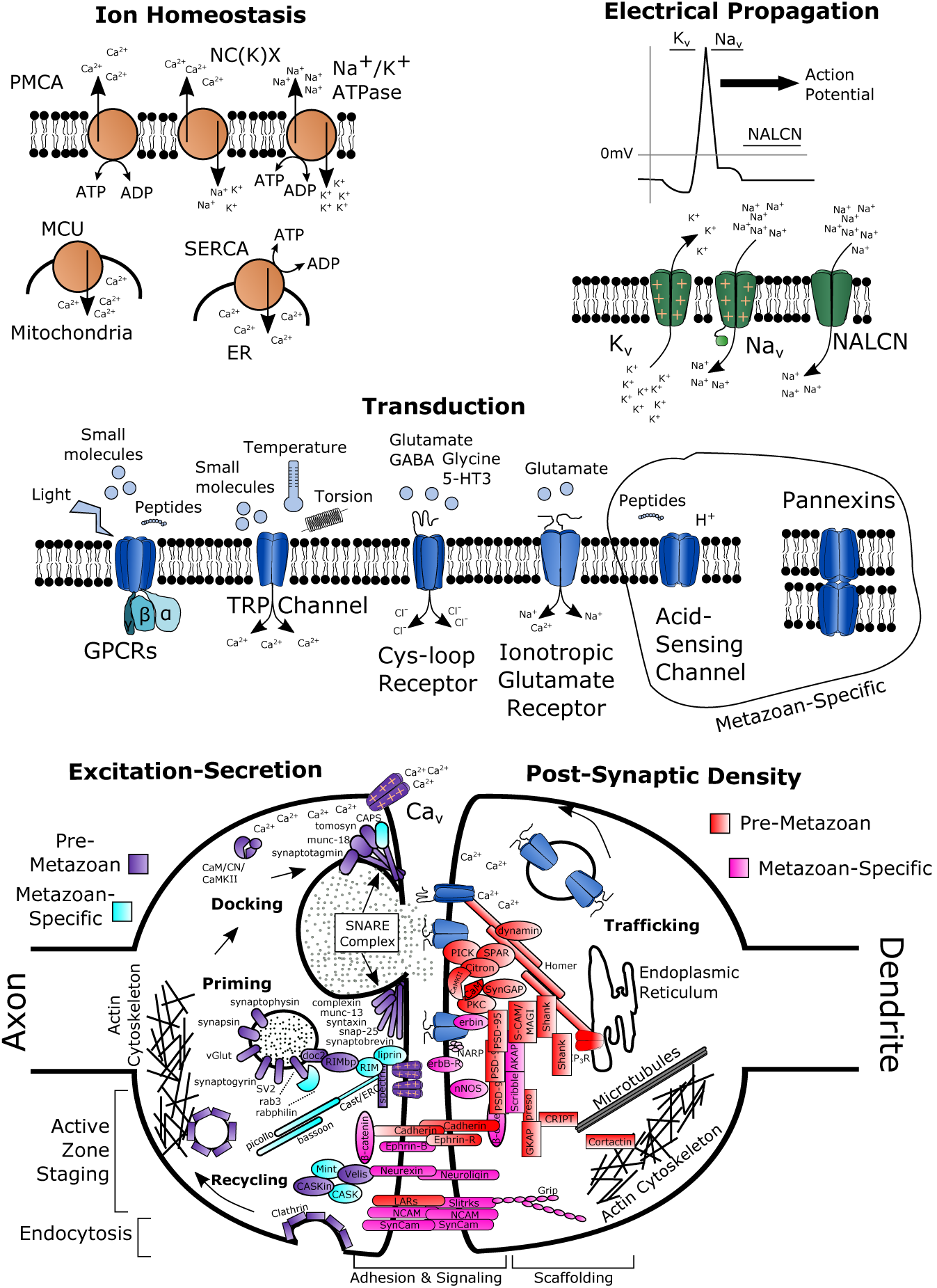
Molecular details of five molecular modules of neurons for which extensive evolutionary information is available. Most channels, pumps and exchangers are evolutionarily ancient, whereas certain sub-modules in synapses are more recent innovations.

### 4.1. Excitation-secretion coupling

Chemical synapses transduce a neurons electrical code into an excreted chemical signal. This happens when the sodium-based action potentials of the axon reach the axon terminus and trigger the opening of Ca_*v*_ channels (Figure 3). The influx of calcium triggers neurotransmitter-filled vesicles to release from their priming area and fuse with the presynaptic membrane. After release, the membrane is then recycled into new vesicles (Figure 3). The proteins involved in docking and in recycling are, for the most part, conserved across eukaryotes [74,75]. The well-known SNARE complex is found across eukaryotes, where it serves a similar function in endomembrane trafficking and exocytosis [76]. Interestingly, SNARE proteins have expanded independently in the major multicellular lineages, suggesting a key role in cell-type differentiation [77]. Clathrin-based endocytosis is also found in all eukaryotes [78]. However, many pre-synaptic proteins involved in scaffolding and in vesicle priming arose only within Metazoa (Figure 3) [74,75].

A nearly complete set of excitation-secretion related proteins are present in choanoflagellates [79], and both sponges and placozoans have a distinct flask-shaped cell-type that is enriched for vesicles and known synaptic genes, and possesses a non-motile cilium, consistent with a role in environmental sensing [80,81]. In the demosponge *Amphimedon queenslandica,* the flask cells have been shown to transduce sensory cues into induction of settlement and metamorphosis in a calcium-dependent manner [82]. Thus these flask cells are probably the best candidates for neuron-like cells in sponges and placozoans, though we caution against interpreting them as proto-neurons.

Several types of molecules are used as neurotransmitters, including amino acids and their derivatives (the biogenic amines), gases (nitric oxide), small peptides, and acetylcholine (Figure 4). Many are used widely in eukaryotes for intercellular communication, but some of the biogenic amines may be present in animals as a result of late horizontal transfer of synthesis enzymes from bacteria [83]. Their deployment in different synapse types across animals is a fascinating and still poorly understood study in evolution. For instance, epinephrine and norepinephrine are important neurotransmitters in vertebrates, but are typically not used at all by protostomes (but see [84]), whereas the opposite is true of octopamine and tyramine (Figure 4). Cnidarians make a similar set of neurotransmitters to vertebrates [85], but *Nematostella* expresses most non-peptide types in the endoderm near the pharynx and testes only peptide transmitters are found in neurons [86]. Ctenophores seem to use a much more restricted set, as glutamate is the only well-validated neurotransmitter [4]. This supports the theory that neurons arose independently in ctenophores and planulozoans, because vertebrate and most protostomes use acetylcholine (AcH) at the neuromuscular junction (NMJ). However, arthropods use glutamate at the NMJ, just like ctenophores [87], and cnidarians probably use neuropeptides [86]. Although sponges do not have true synapses, they use GABA, glutamate, and nitric oxide to coordinate contractions [88]. *Trichoplax* also lacks synapses, but their secretory flask cells label for FMRFamide, suggesting a conserved role in transmission for this peptide class [81].

**Figure 4.**
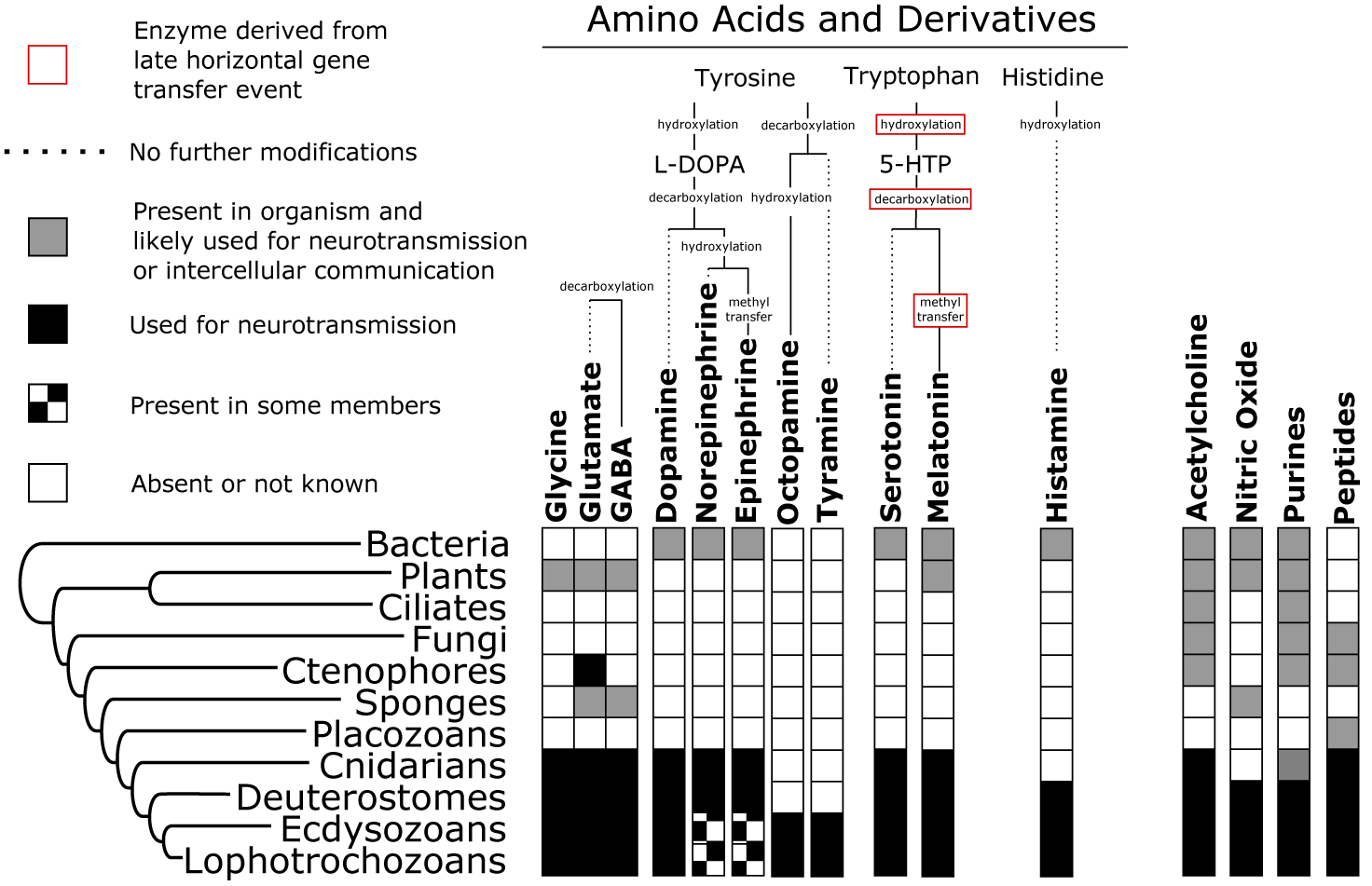
Evolution of neurotransmitter types. Biogenic amines derived from tyrosine, tryptophan, and histadine have a restricted taxon distribution, and may be present in bilaterians+cnidarians as a result of late horizontal gene transfer of key enzymes from bacteria [83]. Other neurotransmitter types are used for intercellular signaling across a wider range of eukaryotes. Presence-absence data is derived from [85,88,121,122].

Calcium-dependent excitation-secretion is therefore an ancient system that has been coopted into synaptic transmission. This cooption involved recruiting neurotransmitters into the vesicle cycle and lineage-specific adaptations for vesicle docking and priming. Much remains to be clarified about how the diverse deployment of neurotransmitters types evolved.

### 4.2. Post-synaptic density

One of the defining histological features of synapses is an electron-dense, disc-shaped structure on the post-synaptic side, called the post-synaptic density (Figure 3). Besides the ion channels that transduce the neurotransmitter signal into post-synaptic potentials, this density includes a large number of scaffolding and signaling proteins that modulate synaptic strength. The proteins of the post-synaptic density can be roughly classified into adhesion and signaling molecules that exist near the post-synaptic membrane, and scaffolding proteins [74]. Many of the adhesion and signaling molecules are newer innovations, particularly those that interact specifically with certain classes of postsynaptic ion channels [75], though much of the signal for a progressive advance in complexity towards mammals probably comes from mammalian-biased sampling of synaptic proteomes [19]. Scaffolding proteins, in contrast, tend to be found outside of Metazoa as well [74,80]. One protein, Homer, has been shown to be ancestrally expressed at the nuclear envelope, based on its expression in choanoflagellates and in mammalian glia [74], highlighting the pleiotropic nature of many of genes in this module.

A nearly complete set of post-synaptic density proteins are found in sponges and in *Trichoplax,* despite the lack of obvious synapses in these organisms [80]. Intriguingly, disc-shaped electron-dense junctions are found in syncytial cell types of hexactinillid sponges [89], *Trichoplax* [81], and colonial choanoflagellates [90]. It is not known what proteins are expressed at these junctions, however, and post-synaptic density genes do not appear to be co-expressed in the sponge *Amphimedon queenslandica* [91]. The flask cells of sponges, which express pre-synaptic genes and have vesicles, also express post-synaptic scaffolding and signaling proteins [80].

### 4.3. Axonogenesis

Neurons are derived from ectoderm in most bilaterians, cnidarians, and ctenophores [92,93]. They can also derive from mesoderm in planarians [94], endoderm in sea urchins [95], and interstitial stem cells in hydrozoan cnidarians [96]. The gene repertoire involved in early neuron specification is largely conserved in bilaterians and cnidarians, but ctenophores are missing some of this core set of neurodevelopment genes, including HOX, Notch, Fibroblast growth factor, and the Hedgehog pathway [5,93]. They do, however, have genes in the Lim-homeobox family (Lhx), which may play a role in neuron development as they do in bilaterians [97]. They also express the Wnt pathway, which is important for body patterning and neurogenesis in cnidarians and bilaterians, but only in adults and only in a very localized area near the mouth [98]. In general, the developmental gene content of ctenophores more closely resembles that of sponges, while the neuron-less *Trichoplax* largely resembles that of cnidarians and bilaterians [5]. Thus the traditional ontogenetic homology criterion cannot be considered full-proof over deep evolutionary time [9].

During development, and after injury, axons must grow and find their correct synaptic targets. The proteins responsible for this targeting include secreted and membrane-bound signals and receptors that have not been studied in an evolutionary framework as well as the other modules. We plotted the evolutionary distribution of some of the best-known families in Figure 5. Most of the axonogenesis proteins have clustered, shared domain repeats [99]. All of the domains shown in Figure 5 are found outside of Metazoa, except for the Semaphorin domain. Proteins with Ig-family domains are well-represented, suggesting a close functional and evolutionary relationship between immunity and axon-guidance. Many these cell adhesion proteins are also found in epithelial junctions, such as the pleated septate junction (non-vertebrates) and tight junctions (vertebrates), and in intermodal septate junctions between myelinating glia and axons. It has therefore been suggested that some synapse types evolved from epithelial precursors [100]. This suggestion is intriguing, as both sponges and placozoans have recently been shown to have occluding *adherens* junctions [101,102]. In fact, both choanoflagellates and the slime mold *Dictyostelium* make electron-dense junctions, the latter of which are known to involve *β*-catenin [90,103].

**Figure 5.**
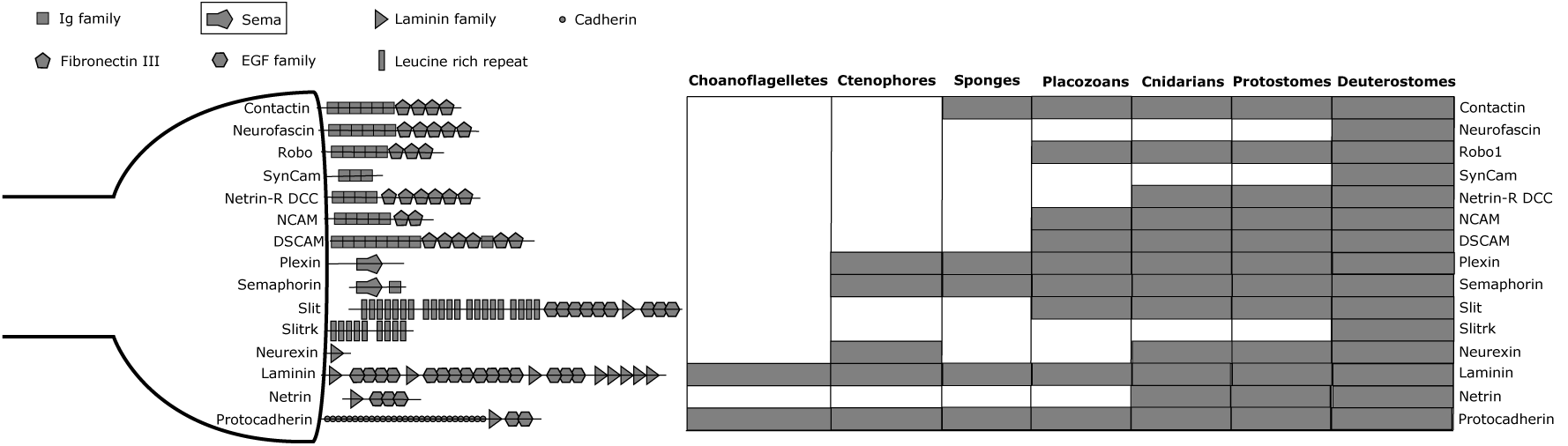
Surface and secreted proteins involved in axonogenesis and synaptic targeting arranged by protein domain composition. Most are made up of multiple domains and have probably evolved by domain shuffling. Many are expressed in other cell types, especially epithelial and immune cells. All domains are found outside of metazoans with the exception of the Semaphorin (SEMA) domain (boxed).

How is the final wiring diagram in the brain specified? In *Drosophila,* the DSCAM gene (Down syndrome cell adhesion molecule) has clustered series of alternate exons that are alternatively spliced to produce an enormous variety of mature protein isoforms. It is thought that this diversity creates a combinatorial code determining the wiring diagram of the fly neural system [104]. In vertebrates, however, DSCAM is not extensively spliced. Instead, vertebrate genomes have clustered arrays of protocadherins, which play a role in dendritic self-avoidance and synaptic pruning [105]. Little is known about these mechanisms outside of current model systems, but, intriguingly, protocadherins have convergently radiated in octopods, where they are also found in clustered genomic repeats and are heavily expressed in the brain [106] a convergent feature of large brains in cephalopods and vertebrates.

Thus the axon guidance module shares many similarities with epithelial and immune junctions. Domain rearrangement and shuffling have probably shaped the distribution of proteins found at synapses. Ctenophores, cnidarians, sponges, placozoans, and choanoflagellates all have undescribed proteins that share domains and overall architecture with animal axon guidance proteins, and much remains to be learned about the deep evolution of synaptic targeting.

## 5. HIERARCHICAL MOLECULAR EVOLUTION

The functional modules described above have different rates and modes of molecular evolution. Some modules are characterized by large gene families and rapid gene turnover (transduction), whereas others have smaller gene families whose size remains fairly stable (ion homeostasis). Functional changes in ion channels, such as those in the propagation or transduction modules, are typically characterized by a few amino acid changes at key sites [51,63,67], whereas adhesion and scaffolding proteins expressed in the axonogenesis and post-synaptic modules may evolve primarily by domain rearrangements and gene fusions [107]. Thus the protein might not even be the relevant homology unit for the latter class of proteins. It even seems possible that whole pathways for neurotransmitter synthesis may have been transferred from bacteria into animals [83]. No gene family or module has an identical taxonomic distribution as neurons and many have quite plastic evolutionary histories.

Currently, the evidence presented for the homology or non-homology of neurons in ctenophores and other animals relies chiefly on comparative genomics data, usually presented as lists of genes or pathways present or not present in the common animal ancestor [12,24]. The considerations above make it clear that not all gene classes are expected to be equally informative for homology considerations. Moreover, a list of genes cannot be interpreted apart from functional context. Proteins function in the hierarchy of molecular processes that make up complex cell types: they form protein complexes, protein complexes function in pathways and subcellular structures, and these structures come together to mediate cell-level phenotypes. At each hierarchical level, evolution at a lower level has a many-to-many mapping to changes at the higher level. Thus it is a non-trivial task to reconstruct the evolutionary dynamics of the higher from the lower level of phenotypes. Doing so becomes more difficult as less is known about intermediate molecular functions, as fewer taxa are sampled across the phylogeny, and as the interior branches of the tree become deeper. Deep neural system evolution exists in the worst part of this space. And yet it is precisely this kind of hierarchical reconstruction that is expected to be most informative for cell-type evolution.

### 5.1. Origins of neuromuscular junctions

One system that might best yield to this sort of analysis is the neuromuscular junction (NMJ). Moroz and colleagues found that the ctenophore NMJ was glutamatergic (glutamate is the neurotransmitter) [4]. Interestingly, this has been used as evidence both for and against the multiple origins hypothesis [12,24]. On one hand vertebrate NMJs are cholinergic (using acetylcholine) and ctenophores have many independently expanded gene families associated with glutamate signaling (although we note that many of the glutamate receptor genes are actually glycine receptors, and the role of glycine signaling in ctenophores is unknown; [52]). On the other hand, glutamatergic synapses and the protein families involved are found across animals. Too little is known to make a determination of homology, but the NMJ makes an interesting case study in how the study of hierarchical molecular evolution is the necessary next step to reconstructing ancient cell types.

The NMJ is a model synapse whose evolution seems likely to reflect that of neurons as a whole; muscles and neurons are always found together in animals, with the sole exception of the parasitic myxozoans, which have lost neurons [108]. Yet, as one looks closer at the molecular composition of NMJs, it becomes apparent that there is considerable evolutionary plasticity. Vertebrates, arthropods, cnidarians, and ctenophores all use different neurotransmitter types at the NMJ, whereas nematodes use the same neurotransmitter as vertebrates (acetylcholine), despite their closer relationship to arthropods (Figure 6). NMJs can also vary considerably in ultrastructure, even among vertebrates [109]. Thus the fact that ctenophore synapses have another distinct morphology [24,73] is not strong evidence of an independent origin. And yet, the distinct pre-synaptic cytomatrix structures in *Drosophila* and *Xenopus* are made up of largely homologous proteins. Some of these homologous proteins (Bruchpilot and ELKS/CAST/ERC) have different compositions of the homologous domains and long non-homologous stretches (Figure 6).

**Figure 6.**
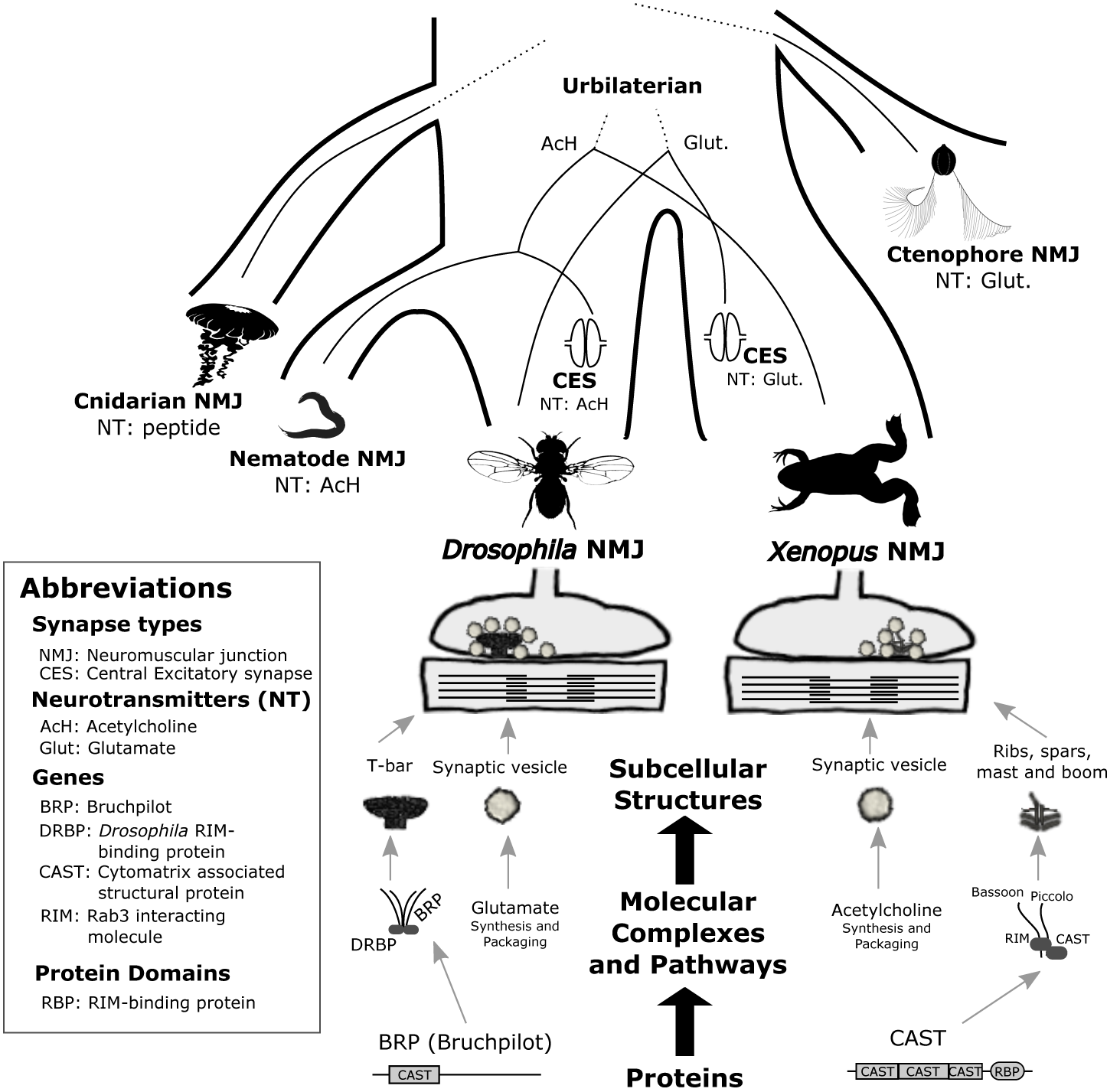
Evolution of the neuromuscular junction (NMJ). There are interesting differences and similarities between two model NMJs, in *Drosophila* and *Xenopus,* when one looks across the hierarchy of molecular organization. Molecular component of *Drosophila* NMJs may be more closely related to excitatory synapses of the brain in vertebrates, and vice versa. Not enough is known about the molecular constituents of ctenophore and cnidarians NMJs to determine their homology with bilaterian cells.

Finally, it has been shown that *Drosophila* NMJs use the same neurotransmitter (glutamate) as vertebrate excitatory synapses in the brain, and also have an overlapping set of cell-adhesion proteins [100]. *Drosophila* central excitatory synapses, inversely, use the same neurotransmitter as vertebrate NMJs (acetylcholine) (Figure 6). Perhaps the last common ancestor of bilaterians had NMJs with multiple transmitter types, as glutamate receptors are expressed at cholinergic NMJs in mammals [110].

Synapses have often been treated as monolithic entities in the evolutionary literature, but much of what is known about the molecular composition of synapses comes from just two synapse types: the excitatory synapses of vertebrates and the *Drosophila* NMJ [111]. But these two glutamatergic synapses have largely overlapping molecular compositions that cannot, without reservation, be extended to other synapse types within one organism [112], let alone to distantly related species. This level of mechanistic detail is known only in a handful of species but is the key to reconstructing the deep evolution of cell-level phenotypes. Thus we are far from making a determination about the homology of synapse types, much less neurons as a whole, across cnidarians, ctenophores, and bilaterians.

All complex cell types are likely to comprise a hierarchy of molecular components with diverse histories in a similar fashion to NMJs, as has been shown to be the case with muscle types [113,114]. It should be acknowledged that all possible patterns of co-evolution across levels are not only possible but plausible over even modest evolutionary divergences. Thus, molecular similarities between similar phenotypes cannot be unambiguously interpreted as signs of homology, and molecular differences are not necessarily signs of convergence. Proper interpretation will only be possible when the intrinsic rates of mechanism-phenotype coevolution are known for particular phenotype-mechanism pairs.

Will such rates ever be known for phenotypes of interest? Convergent evolution and systems drift of neural systems on higher organizational levels is well known [115]. Famous examples of convergence include the multiple independent origins of myelination and of giant axons for rapid electrical signal propagation [28]; of electric organs in two lineages of teleosts and in elasmobranchs [116]; of higher-order brain anatomy in vertebrates, arthropods, and molluscs [11,117,118]; and of chambered eyes in vertebrates and cephalopods [119]. Systems drift is less well-studied, but is clearly widespread, especially on the level of specific brain circuits [118,120]. Thus, comparative studies of neural system evolution in more closely related species can lead the way.

## 6. CONCLUSION

Neural systems are spectacularly complex phenotypes. Comparative genomics has opened the door to evolutionary neuroscience across the most divergent animal taxa. Future studies will take advantage of these advances by detailing the molecular composition and function of neural systems in non-traditional model organisms and by developing comparative methods that can deal with the intrinsic hierarchy of molecular systems.

## ACKNOWLEDGMENTS

We would like to acknowledge Dr. Rebecca Young for helpful discussions. B.J.L. was funded by NIH fellowship 1F32GM112504-01A1. H.A.H acknowledges grants NSF IOS-1354942, NSF IOS-1501704, and Alfred P. Sloan Foundation (BR-4900)

## LITERATURE CITED

1. van Leewenhoeck, A. 1677. Observations, Communicated to the Publisher by Mr. Antony van Leewenhoeck, in a Dutch Letter of the 9th of Octob. 1676. Here Englishd: concerning Little Animals by Him Observed in Rain–Well–Sea. and Snow Water; as Also in Water Wherein Pepper Had Lain Infused. Philos. Trans. R. Soc. Lond. 12, 821-831.

2. Abedin, M., and King, N. 2010. Diverse evolutionary paths to cell adhesion. Trends Cell Biol. 20, 734–742.

3. Babonis, L.S., and Martindale, M.Q. 2017. Phylogenetic evidence for the modular evolution of metazoan signalling pathways. Phil Trans R Soc B 372, 20150477.

4. Moroz, L.L., Kocot, K.M., Citarella, M.R., Dosung, S., Norekian, T.P., Povolotskaya, I.S., Grigorenko, A.P., Dailey, C., Berezikov, E., Buckley, K.M., et al. 2014. The ctenophore genome and the evolutionary origins of neural systems. Nature 510, 109–114.

5. Ryan, J.F., Pang, K., Schnitzler, C.E., Nguyen, A.D., Moreland, R.T., Simmons, D.K., Koch, B.J., Francis, W.R., Havlak, P., Smith, S.A., et al. 2013. The genome of the ctenophore *Mnemiopsis leidyi* and its implications for cell type evolution. Science 342, 1242592.

6. Srivastava, M., Begovic, E., Chapman, J., Putnam, N.H., Hellsten, U., Kawashima, T., Kuo, A., Mitros, T., Salamov, A., Carpenter, M.L., et al. 2008. The *Trichoplax* genome and the nature of placozoans. Nature 454, 955–960.

7. Srivastava, M., Simakov, O., Chapman, J., Fahey, B., Gauthier, M.E.A., Mitros, T., Richards, G.S., Conaco, C., Dacre, M., Hellsten, U., et al. 2010. The *Amphimedon queenslandica* genome and the evolution of animal complexity. Nature 466, 720–726.

8. Dunn, C.W., Giribet, G., Edgecombe, G.D., and Hejnol, A. 2014. Animal phylogeny and its evolutionary implications. Annu. Rev. Ecol. Evol. Syst. 45, 371–395.

9. Hejnol, A., and Lowe, C.J. 2015. Embracing the comparative approach: how robust phylogenies and broader developmental sampling impacts the understanding of nervous system evolution. Phil Trans R Soc B 370, 20150045.

10. Jékely, G., Paps, J., and Nielsen, C. 2015. The phylogenetic position of ctenophores and the origins of nervous systems. EvoDevo 6

11. Moroz, L.L. 2009. On the independent origins of complex brains and neurons. Brain. Behav. Evol. 74, 177–190.

12. Ryan, J.F. 2014. Did the ctenophore nervous system evolve independently? Zoology 117, 225–226.

13. Bishop, G.H. 1956. Natural history of the nerve impulse. Physiol. Rev. 36, 376–399.

14. Mackie, G.O. 1990. The elementary nervous system revisited. Am. Zool. 30, 907–920.

15. Pisani, D., Pett, W., Dohrmann, M., Feuda, R., Rota-Stabelli, O., Philippe, H., Lartillot, N., and Wörheide, G. 2015. Genomic data do not support comb jellies as the sister group to all other animals. Proc. Natl. Acad. Sci., 201518127.

16. Arcila, D., Ortí, G., Vari, R., Armbruster, J.W., Stiassny, M.L.J., Ko, K.D., Sabaj, M.H., Lundberg, J., Revell, L.J., and Betancur,R, R. 2017. Genome–wide interrogation advances resolution of recalcitrant groups in the tree of life. Nat. Ecol. Evol. 1, 0020.

17. Dunn, C.W., Hejnol, A., Matus, D.Q., Pang, K., Browne, W.E., Smith, S.A., Seaver, E., Rouse, G.W., Obst, M., Edgecombe, G.D., et al. 2008. Broad phylogenomic sampling improves resolution of the animal tree of life. Nature 452, 745–749.

18. Whelan, N.V., Kocot, K.M., Moroz, L.L., and Halanych, K.M. 2015. Error, signal, and the placement of Ctenophora sister to all other animals. Proc. Natl. Acad. Sci. 112, 5773–5778.

19. Liebeskind, B.J., Hillis, D.M., Zakon, H.H., and Hofmann, H.A. 2016. Complex Homology and the Evolution of Nervous Systems. Trends Ecol. Evol. 31, 127–135.

20. Cunningham, J.A., Liu, A.G., Bengtson, S., and Donoghue, P.C.J. 2016. The origin of animals: Can molecular clocks and the fossil record be reconciled. BioEssays 39, 1600120.

21. dos Reis, M., Thawornwattana, Y., Angelis, K., Telford, M.J., Donoghue, P.C.J., and Yang, Z. 2015. Uncertainty in the Timing of Origin of Animals and the Limits of Precision in Molecular Timescales. Curr. Biol. 25, 2939–2950.

22. True, J.R., and Haag, E.S. 2001. Developmental system drift and flexibility in evolutionary trajectories. Evol. Dev. 3, 109–119.

23. Shubin, N., Tabin, C., and Carroll, S. 1997. Fossils, genes and the evolution of animal limbs. Nature 388, 639–648.

24. Moroz, L.L., and Kohn, A.B. 2016. Independent origins of neurons and synapses: insights from ctenophores. Phil Trans R Soc B 371, 20150041.

25. Edwards, D.H., Heitler, W.J., and Krasne, F.B. 1999. Fifty years of a command neuron: the neurobiology of escape behavior in the crayfish. Trends Neurosci. 22, 153–161.

26. Mackie, G.O. 1989. Evolution of cnidarian giant axons. Evolution of the First Nervous Systems NATO ASI Series., P. A. V. Anderson, ed. Springer US, pp. 395–407.

27. Mackie, G.O., Mills, C.E., and Singla, C.L. 1992. Giant axons and escape swimming in *Euplokamis dunlapae* Ctenophora: Cydippida. Biol Bull 182, 248–256.

28. Castelfranco, A.M., and Hartline, D.K. 2016. Evolution of rapid nerve conduction. Brain Res. 1641, Part A, 11–33.

29. Laughlin, S.B., Steveninck, R.R. de R. van, and Anderson, J.C. 1998. The metabolic cost of neural information. Nat. Neurosci. 1, 36–41.

30. Thever, M.D., and Saier, M.H. 2009. Bioinformatic characterization of P–Type ATPases encoded within the fully sequenced genomes of 26 eukaryotes. J. Membr. Biol. 229, 115–130.

31. Barrero–Gil, J., Garciadeblás, B., and Benito, B. 2005. Sodium, potassium–ATPases in algae and oomycetes. J. Bioenerg. Biomembr. 37, 269–278.

32. Cai, X., and Clapham, D.E. 2012. Ancestral Ca^2^ + signaling machinery in early animal and fungal evolution. Mol. Biol. Evol. 29, 91–100.

33. Cai, X., and Lytton, J. 2004. The cation/Ca^2^ + exchanger superfamily: phylogenetic analysis and structural implications. Mol. Biol. Evol. 21, 1692–1703.

34. Jaiteh, M., Taly, A., and Hénin, J. 2016. Evolution of pentameric ligand–gated ion channels: Pro–loop receptors. PLOS ONE 11, e0151934.

35. Liebeskind, B.J., Hillis, D.M., and Zakon, H.H. 2015. Convergence of ion channel genome content in early animal evolution. Proc. Natl. Acad. Sci. 112, E846–E851.

36. Dalton, R.P., and Lomvardas, S. 2015. Chemosensory receptor specificity and regulation. Annu. Rev. Neurosci. 38, 331–349.

37. Touhara, K., and Vosshall, L.B. 2009. Sensing odorants and pheromones with chemosensory receptors. Annu. Rev. Physiol. 71, 307–332.

38. Niimura, Y. 2012. Olfactory receptor multigene family in vertebrates: from the viewpoint of evolutionarygenomics. Curr. Genomics 13, 103–114.

39. Nei, M., and Rooney, A.P. 2005. Concerted and Birth–and–Death Evolution of Multigene Families. Annu. Rev. Genet. 39, 121–152.

40. Sakarya, O., Kosik, K.S., and Oakley, T.H. 2008. Reconstructing ancestral genome content based on symmetrical best alignments and Dollo parsimony. Bioinformatics 24, 606–612.

41. Liegertová, M., Pergner, J., Kozmiková, I., Fabian, P., Pombinho, A.R., Strnad, H., Paces, J., Vlcek, C., Bartunek, P., and Kozmik, Z. 2015. Cubozoan genome illuminates functional diversification of opsins and photoreceptor evolution. Sci. Rep. 5, 11885

42. Rivera, A.S., Ozturk, N., Fahey, B., Plachetzki, D.C., Degnan, B.M., Sancar, A., and Oakley, T.H. 2012. Blue-light-receptive cryptochrome is expressed in a sponge eye lacking neurons and opsin. J. Exp. Biol. 215, 1278–1286.

43. Silbering, A.F., and Benton, R. 2010. Ionotropic and metabotropic mechanisms in chemoreception: chance or design? EMBO Rep. 11, 173–179.

44. Croset, V., Rytz, R., Cummins, S.F., Budd, A., Brawand, D., Kaessmann, H., Gibson, T.J., and Benton, R. 2010. Ancient protostome origin of chemosensory ionotropic glutamate receptors and the evolution of insect taste and olfaction. PLOS Genet. 6, e1001064.

45. Saina, M., Busengdal, H., Sinigaglia, C., Petrone, L., Oliveri, P., Rentzsch, F., and Benton, R. 2015. A cnidarian homologue of an insect gustatory receptor functions in developmental body patterning. Nat. Commun. 6, 6243.

46. Julius, D., and Nathans, J. 2012. Signaling by sensory receptors. Cold Spring Harb. Perspect. Biol. 4, a005991.

47. Bianchi, L., and Driscoll, M. 2002. Protons at the gate: DEG/ENaC ion channels help us feel and remember. Neuron 34, 337–340.

48. Assmann, M., Kuhn, A., Dürrnagel, S., Holstein, T.W., and Gründer, S. 2014. The comprehensive analysis of DEG/ENaC subunits in Hydra reveals a large v ariety of peptide–gated channels, potentially involved in neuromuscular transmission. BMC Biol. 12, 84.

49. Chiu, J., DeSalle, R., Lam, H.M., Meisel, L., and Coruzzi, G. 1999. Molecular evolution of glutamate receptors: a primitive signaling mechanism that existed before plants and animals diverged. Mol. Biol. Evol. 16, 826–838.

50. Kehoe, J., Buldakova, S., Acher, F., Dent, J., Bregestovski, P., and Bradley, J. 2009. Aplysia cys–loop glutamate–gated chloride channels reveal convergent evolution of ligand specificity. J. Mol. Evol. 69, 125–141.

51. Lynagh, T., Beech, R.N., Lalande, M.J., Keller, K., Cromer, B.A., Wolstenholme, A.J., and Laube, B. 2015. Molecular basis for convergent evolution of glutamate recognition by pentameric ligand–gated ion channels. Sci. Rep. 5, 8558.

52. Alberstein, R., Grey, R., Zimmet, A., Simmons, D.K., and Mayer, M.L. 2015. Glycine activated ion channel subunits encoded by ctenophore glutamate receptor genes. Proc. Natl. Acad. Sci. 112, E6048–6057.

53. Thornton, J.W. 2001. Evolution of vertebrate steroid receptors from an ancestral estrogen receptor by ligand exploitation and serial genome expansions. Proc. Natl. Acad. Sci. 98, 5671–5676.

54. Simons, P. j. 1981. The Role of Electricity in Plant Movements. New Phytol. 87, 11–37.

55. Oami, K., Naitoh, Y., and Sibaoka, T. 1995. Voltage-gated ion conductances corresponding to regenerative positive and negative spikes in the dinoflagellateNoctiluca miliaris. J. Comp. Physiol. A 176, 625–633.

56. Febvre–chevalier, C., Bilbaut, A., Bone, Q., and Febvre, J. 1986. Sodium–calcium action potential associated with contraction in the heliozoan *Actinocoryne contractilis*. J. Exp. Biol. 122, 177–192.

57. Saimi, Y., and Kung, C. 1987. Behavioral Genetics of Paramecium. Annu. Rev. Genet. 21, 47–65.

58. Prindle, A., Liu, J., Asally, M., Ly, S., Garcia–Ojalvo, J., and Süel, G.M. 2015. Ion channels enable electrical communication in bacterial communities. Nature 527, 59-63

59. Hille, B. 2001. Ion Channels of Excitable Membranes 3^rd^ Edition. Sunderland, MA: Sinauer Associates.

60. Lishko, P.V., Kirichok, Y., Ren, D., Navarro, B., Chung, J.-J., and Clapham, D.E. 2012. The control of male fertility by spermatozoan ion Channels. Annu. Rev. Physiol. 74, 453–475.

61. Ren, D. 2011. Sodium leak channels in neuronal excitability and rhythmic behaviors. Neuron 72, 899–911.

62. Gur Barzilai, M., Reitzel, A.M., Kraus, J.E.M., Gordon, D., Technau, U., Gurevitz, M., and Moran, Y. 2012. Convergent evolution of sodium ion selectivity in metazoan neuronal signaling. Cell Rep. 2, 242–248.

63. Liebeskind, B.J., Hillis, D.M., and Zakon, H.H. 2011. Evolution of sodium channels predates the origin of nervous systems in animals. Proc. Natl. Acad. Sci. 108, 9154–9159.

64. Liebeskind, B.J., Hillis, D.M., and Zakon, H.H. 2012. Phylogeny unites animal sodium leak channels with fungal calcium channels in an ancient, voltage–insensitive clade. Mol. Biol. Evol. 29, 3613–3616.

65. Senatore, A., Monteil, A., van Minnen, J., Smit, A.B., and Spafford, J.D. 2013. NALCN ion channels have alternative selectivity filters resembling calcium channels or sodium channels. PLOS ONE 8, e55088.

66. Li, X., Liu, H., Luo, J.C., Rhodes, S.A., Trigg, L.M., Rossum, D.B. van, Anishkin, A., Diatta, F.H., Sassic, J.K., Simmons, D.K., et al. 2015. Major diversification of voltage–gated K^+^ channels occurred in ancestral parahoxozoans. Proc. Natl. Acad. Sci. 112, E1010–E1019.

67. Martinson, A.S., Rossum, D.B. van, Diatta, F.H., Layden, M.J., Rhodes, S.A., Martindale, M.Q., and Jegla, T. 2014. Functional evolution of Erg potassium channel gating reveals an ancient origin for IKr. Proc. Natl. Acad. Sci. 111, 5712–5717

68. Bilbaut, A., Hernandez–Nicaise, M.L., and Meech, R.W. 1989. Ionic Currents in Ctenophore Muscle Cells. Evolution of the First Nervous Systems NATO ASI Series., P. A. V. Anderson, ed. Springer US, pp. 299–314.

69. Leys, S., Mackie, G., and Meech, R. 1999. Impulse conduction in a sponge. J Exp Biol 202, 1139–1150.

70. Pereda, A.E. 2014. Electrical synapses and their functional interactions with chemical synapses. Nat. Rev. Neurosci. 15, 250–263.

71. Haas, J.S., Zavala, B., and Landisman, C.E. 2011. Activity–dependent long–term depression of electrical synapses. Science 334, 389–393.

72. Anderson, P.A. 1985. Physiology of a bidirectional, excitatory, chemical synapse. J. Neurophysiol. 53, 821–835.

73. Hernandez–Nicaise, M.L. 1973. The nervous system of ctenophores III. Ultrastructure of synapses. J. Neurocytol. 2, 249–263.

74. Burkhardt, P., Grønborg, M., McDonald, K., Sulur, T., Wang, Q., and King, N. 2014. Evolutionary insights into premetazoan functions of the neuronal protein Homer. Mol. Biol. Evol.. 31, 2342-2355

75. Emes, R.D., and Grant, S.G.N. 2012. Evolution of synapse complexity and diversity. Annu. Rev. Neurosci. 35, 111–131.

76. Kloepper, T.H., Kienle, C.N., and Fasshauer, D. 2007. An elaborate classification of SNARE proteins sheds light on the conservation of the eukaryotic endomembrane system. Mol. Biol. Cell. 18, 3463–3471.

77. Richter, D.J., and King, N. 2013. The genomic and cellular foundations of animal origins. Annu. Rev. Genet. 47, 509–537.

78. Becker, B., and Melkonian, M. 1996. The secretory pathway of protists: spatial and functional organization and evolution. Microbiol. Rev. 60, 697–721.

79. Burkhardt, P., Stegmann, C.M., Cooper, B., Kloepper, T.H., Imig, C., Varoqueaux, F., Wahl, M.C., and Fasshauer, D. 2011. Primordial neurosecretory apparatus identified in the choanoflagellate *Monosiga brevicollis*. Proc. Natl. Acad. Sci. 108, 15264–15269.

80. Sakarya, O., Armstrong, K.A., Adamska, M., Adamski, M., Wang, I.F., Tidor, B., Degnan, B.M., Oakley, T.H., and Kosik, K.S. 2007. A Post-Synaptic Scaffold at the Origin of the Animal Kingdom. PLOS ONE 2, e506.

81. Smith, C.L., Varoqueaux, F., Kittelmann, M., Azzam, R.N., Cooper, B., Winters, C.A., Eitel, M., Fasshauer, D., and Reese, T.S. 2014. Novel cell types, neurosecretory cells, and body plan of the early–diverging metazoan Trichoplax adhaerens. Curr. Biol. 24, 1565–1572.

82. Nakanishi, N., Stoupin, D., Degnan, S.M., and Degnan, B.M. 2015. Sensory flask cells in sponge larvae regulate metamorphosis *via* calcium signaling. Integr. Comp. Biol., icv014.

83. Iyer, L.M., Aravind, L., Coon, S.L., Klein, D.C., and Koonin, E.V. 2004. Evolution of cell–cell signaling in animals: did late horizontal gene transfer from bacteria have a role? Trends Genet. 20, 292–299.

84. Bauknecht, P., and Jékely, G. 2017. Ancient coexistence of norepinephrine, tyramine, and octopamine signaling in bilaterians. BMC Biol. 15, 6.

85. Kass–Simon, G., and Pierobon, P. 2007. Cnidarian chemical neurotransmission, an updated overview. Comp. Biochem. Physiol. A. Mol. Integr. Physiol. 146, 9–25.

86. Oren, M., Brikner, I., Appelbaum, L., and Levy, O. 2014. Fast neurotransmission related genes are expressed in non nervous endoderm in the sea anemone *Nematostella vectensis*. PLOS ONE 9, e93832.

87. Jan, L.Y., and Jan, Y.N. 1976. L-glutamate as an excitatory transmitter at the *Drosophila* larval neuromuscular junction. J. Physiol. 262, 215–236.

88. Elliott, G.R.D., and Leys, S.P. 2010. Evidence for glutamate, GABA and NO in coordinating behaviour in the sponge, *Ephydatia muelleri* Demospongiae, Spongillidae. J. Exp. Biol. 213, 2310–2321.

89. Leys, S.P. 2003. The significance of syncytial tissues for the position of the Hexactinellida in the Metazoa. Integr. Comp. Biol. 43, 19–27.

90. Dayel, M.J., Alegado, R.A., Fairclough, S.R., Levin, T.C., Nichols, S.A., McDonald, K., and King, N. 2011. Cell differentiation and morphogenesis in the colony–forming choanoflagellate *Salpingoeca rosetta*. Dev. Biol. 357, 73–82.

91. Conaco, C., Bassett, D.S., Zhou, H., Arcila, M.L., Degnan, S.M., Degnan, B.M., and Kosik, K.S. 2012. Functionalization of a protosynaptic gene expression network. Proc. Natl. Acad. Sci. 109, 10612–10618.

92. Hartenstein, V., and Stollewerk, A. 2015. The Evolution of early neurogenesis. Dev. Cell. 32, 390–407.

93. Martindale, M.Q., and Henry, J.Q. 2015. Ctenophora. Evolutionary Developmental Biology of Invertebrates 1, A. Wanninger, ed. Vienna: Springer, pp. 179–201.

94. Rink, J.C. 2013. Stem cell systems and regeneration in planaria. Dev. Genes Evol. 223, 67–84.

95. Wei, Z., Angerer, R.C., and Angerer, L.M. 2011. Direct development of neurons within foregut endoderm of sea urchin embryos. Proc. Natl. Acad. Sci. 108, 9143–9147.

96. Bode, H.R. 1996. The interstitial cell lineage of hydra: a stem cell system that arose early in evolution. J. Cell Sci. 109, 1155–1164.

97. Simmons, D.K., Pang, K., and Martindale, M.Q. 2012. Lim homeobox genes in the Ctenophore Mnemiopsis leidyi: the evolution of neural cell type specification. EvoDevo 3, 2.

98. Jager, M., Chiori, R., Alié, A., Dayraud, C., Quéinnec, E., and Manuel, M. 2011. New insights on ctenophore neural anatomy: Immunofluorescence study in *Pleurobrachia pileus* Mller, 1776. J. Exp. Zoolog. B Mol. Dev. Evol. 316B, 171–187.

99. de Wit, J., and Ghosh, A. 2016. Specification of synaptic connectivity by cell surface interactions. Nat. Rev. Neurosci. 17, 4–4.

100. Harden, N., Wang, S.J.H., and Krieger, C. 2016. Making the connection – shared molecular machinery and evolutionary links underlie the formation and plasticity of occluding junctions and synapses. J Cell Sci 129, 3067–3076.

101. Adams, E.D.M., Goss, G.G., and Leys, S.P. 2010. Freshwater sponges have functional, sealing epithelia with high transepithelial resistance and negative transepithelial potential. PLOS ONE 5.

102. Smith, C.L., and Reese, T.S. 2016. Adherens junctions modulate diffusion between epithelial cells in *Trichoplax adhaerens*. Biol. Bull. 231, 216–224.

103. Grimson, M.J., Coates, J.C., Reynolds, J.P., Shipman, M., Blanton, R.L., and Harwood, A.J. 2000. Adherens junctions and -catenin-mediated cell signalling in a non-metazoan organism. Nature 408, 727–731.

104. Schmucker, D., Clemens, J.C., Shu, H., Worby, C.A., Xiao, J., Muda, M., Dixon, J.E., and Zipursky, S.L. 2000. Drosophila DSCAM is an axon guidance receptor exhibiting extraordinary molecular diversity. Cell 101, 671–684.

105. Kostadinov, D., and Sanes, J.R. 2015. Protocadherin–dependent dendritic self-avoidance regulates neural connectivity and circuit function. eLife 4, e08964.

106. Albertin, C.B., Simakov, O., Mitros, T., Wang, Z.Y., Pungor, J.R., Edsinger–Gonzalez, E., Brenner, S., Ragsdale, C.W., and Rokhsar, D.S. 2015. The octopus genome and the evolution of cephalopod neural and morphological novelties. Nature 524, 220–224.

107. Schmitz, F., Königstorfer, A., and Südhof, T.C. 2000. RIBEYE, a component of synaptic ribbons: A proteins journey through evolution provides insight into synaptic ribbon function. Neuron 28, 857–872.

108. Gruhl, A., and Okamura, B. 2015. Tissue Characteristics and Development in Myxozoa. Myxozoan Evolution, Ecology and Development, B. Okamura, A.. Gruhl, and J. L.. Bartholomew, eds. New York: Springer International Publishing, pp. 155–174.

109. Ackermann, F., Waites, C.L., and Garner, C.C. 2015. Presynaptic active zones in invertebrates and vertebrates. EMBO Rep. 16, 923–938.

110. Mays, T.A., Sanford, J.L., Hanada, T., Chishti, A.H., and Rafael–Fortney, J.A. 2009. Glutamate receptors localize postsynaptically at neuromuscular junctions in mice. Muscle Nerve 39, 343–349.

111. Husi, H., Ward, M.A., Choudhary, J.S., Blackstock, W.P., and Grant, S.G.N. 2000. Proteomic analysis of NMDA receptor-adhesion protein signaling complexes. Nat. Neurosci. 3, 661–669.

112. ORourke, N.A., Weiler, N.C., Micheva, K.D., and Smith, S.J. 2012. Deep molecular diversity of mammalian synapses: why it matters and how to measure it. Nat. Rev. Neurosci. 13, 365–379.

113. Brunet, T., Fischer, A.H., Steinmetz, P.R., Lauri, A., Bertucci, P., and Arendt, D. 2016. The evolutionary origin of bilaterian smooth and striated myocytes. eLife 5, e19607.

114. Steinmetz, P.R.H., Kraus, J.E.M., Larroux, C., U. Hammel, J., Amon-Hassenzahl, A., Houliston, E., Worheide, G., Nickel, M., Degnan, B.M., and Technau, U. 2012. Independent evolution of striated muscles in cnidarians and bilaterians. Nature 487, 231–234.

115. Nishikawa, K.C. 2002. Evolutionary convergence in nervous systems: insights from comparative phylogenetic studies. Brain. Behav. Evol. 59, 240.

116. Thompson, A., Vo, D., Comfort, C., and Zakon, H.H. 2014. Expression evolution facilitated the convergent neofunctionalization of a sodium channel Gene. Mol. Biol. Evol. 31, 1941–1955.

117. Farris, S.M. 2008. Evolutionary convergence of higher brain centers spanning the protostome– deuterostome boundary. Brain. Behav. Evol. 72, 106–122.

118. Katz, P.S. 2016. Phylogenetic plasticity in the evolution of molluscan neural circuits. Curr. Opin. Neurobiol. 41, 8–16.

119. Gehring, W.J. 2005. New perspectives on eye development and the evolution of eyes and photoreceptors. J. Hered. 96, 171–184.

120. OConnell, L.A., and Hofmann, H.A. 2011. The vertebrate mesolimbic reward system and social behavior network: a comparative synthesis. J. Comp. Neurol. 519, 3599–3639.

121. Roshchina, V.V. 2010. Evolutionary considerations of neurotransmitters in microbial, plant, and animal cells. Microbial Endocrinology, M. Lyte and P. P. E. Freestone, New York: Springer, pp. 17–52.

122. Ruggieri, R.D., Pierobon, P., and Kass-Simon, G. 2004. Pacemaker activity in hydra is modulated by glycine receptor ligands. Comp. Biochem. Physiol. A. Mol. Integr. Physiol. 138, 193–202.

